# Stable 3D Head Direction Signals in the Primary Visual Cortex

**DOI:** 10.1101/2020.09.04.283762

**Authors:** Grigori Guitchounts, William Lotter, Joel Dapello, David Cox

**Affiliations:** Center for Brain Science, Harvard University, Cambridge, Massachusetts 02138, USA; Department of Molecular and Cellular Biology, Harvard University, Cambridge, Massachusetts, 02138, USA; Program in Neuroscience, Harvard University, Cambridge, Massachusetts 02138, USA; School of Engineering and Applied Sciences, Harvard University, Cambridge, Massachusetts 02139, USA

**Keywords:** Head Direction, Visual Cortex, Predictive Coding, Movement, Naturalistic Behavior, Navigation

## Abstract

Visual signals influence the brain’s computation of spatial position and orientation. Accordingly, the primary visual cortex (V1) is extensively interconnected with areas involved in computing head direction (HD) information. Predictive coding theories posit that higher cortical areas send sensory or motor predictions to lower areas, but whether this includes cognitive variables like the HD signal—and whether HD information is present in V1—is unknown. Here we show that V1 encodes the yaw, roll, and pitch of the head in freely behaving rats, either in the presence or absence of visual cues. HD tuning was modulated by lighting and movement state, but was stable on a population level for over a week. These results demonstrate the presence of a critical spatial orientation signal in a primary cortical sensory area and support predictive coding theories of brain function.

## Introduction

Navigation is a core cognitive process critical for survival. The mammalian brain’s navigation systems contain specialized cells whose dynamics reflect two key properties of navigation: a sense of location—via place, grid, and border cells—and heading direction or spatial orientation^1,2^. Head direction (HD) cells, which signal spatial orientation, are found in a number of species^3–9^ and brain areas; in rodents, these include the thalamus, hippocampal formation, and the neocortex^10–16^. HD cells rely on vestibular signals relayed via brainstem nuclei ^10^, and are modulated by visual landmark signals sent from visual areas to the retrosplenial cortex and subiculum^11^. While the influence of visual cues on navigational variables has been studied extensively, it remains largely unknown what effect, if any, navigational variables–a sense of location and heading direction or orientation in space–have on visual representation.

While visual cortical areas have traditionally been thought of as processing centers that transform and analyze incoming sensory information^17^, there is a growing awareness that feedback and modulatory inputs from other modalities have profound effects on visual cortex dynamics^18–21^. Neurons in primary visual cortex (V1) have been shown to signal running speed^22^, increase the gain on visual stimulus during locomotion^18^, and even to signal the direction of movement of the head in the presence or absence of visual input ^23,24^.

Importantly, V1 neurons have also been shown to encode a subjective sense of position in physical space, reminiscent of hippocampal place cells ^25–28^. While the functional purpose and circuit mechanisms for place representation in V1 are not clear, some have suggested that these signals represent the brain’s internal model of the world, as part of a predictive coding framework^29^. Nevertheless, an outstanding question is what functional purpose, if any, navigational signals such as the sense of location or head direction serve for sensory processing.

A major source of non-visual input to V1 comes from the retrosplenial cortex (RSC), a multimodal area known to contain HD cells ^19,30–32^. RSC inputs to V1 have been shown to reflect movement ^20^, but the full functional consequences of these inputs on V1 are not well understood, and the extent to which V1 activity represents head direction is an open question. To test whether V1 dynamics reflect HD, we recorded neuronal activity in V1 of freely behaving rats in an open arena home-cage. Half of the sessions were recorded in the dark in order to assess non-visual HD representations in V1. Many individual V1 neurons were tuned to the yaw, roll, or pitch of the head, in either the dark or the light. These features, which constitute the orientation of the head in 3D space, could be decoded well above chance levels with simple linear models from either a mesoscopic multiunit signal or single-unit firing rates in V1. While individual V1 neurons exhibited variation in tuning to HD based on the lighting condition or movement state, the population-level HD signal was stable for over one week. Altogether, these results indicate that the HD signal extends to a primary sensory cortical area, and suggest that spatial orientation signals are more widespread than previously known.

## Results

### V1 Neurons are Tuned to Head Direction in 3D

To investigate whether V1 encodes HD information, we recorded neuronal activity using tetrode arrays targeting layer 2/3 of rat V1 while the animals behaved freely in a home-cage arena (Figures 1a,b, S1a-c). Movements were captured using a head-mounted inertial measurement unit (IMU) that combined sensor readings from an accelerometer, gyroscope, and magnetometer (Figure S1d-f). We focused our analysis on the Euler angles—yaw (a.k.a. azimuth), roll, and pitch—which allowed analysis of 3D direction of the head in allocentric room coordinates. Recordings were performed continuously, 24/7, with pseudo-randomized dark or light epochs that overrode the animals’ natural dark-light cycles to control for possible circadian effects. The continuous recordings were split into ∼ 2-hour dark or light sessions for analysis purposes, which allowed us to examine HD signals in the absence or presence of visual inputs, respectively. Precautions were taken to ensure light levels in the behavioral box in the dark were lower than what is likely required to orient in space using visual landmarks (see Materials and Methods), signifying that any HD signals in V1 in the dark correspond to an estimate based on nonvisual signals (e.g. vestibular, somatosensory, auditory, or olfactory).

**Figure 1:**
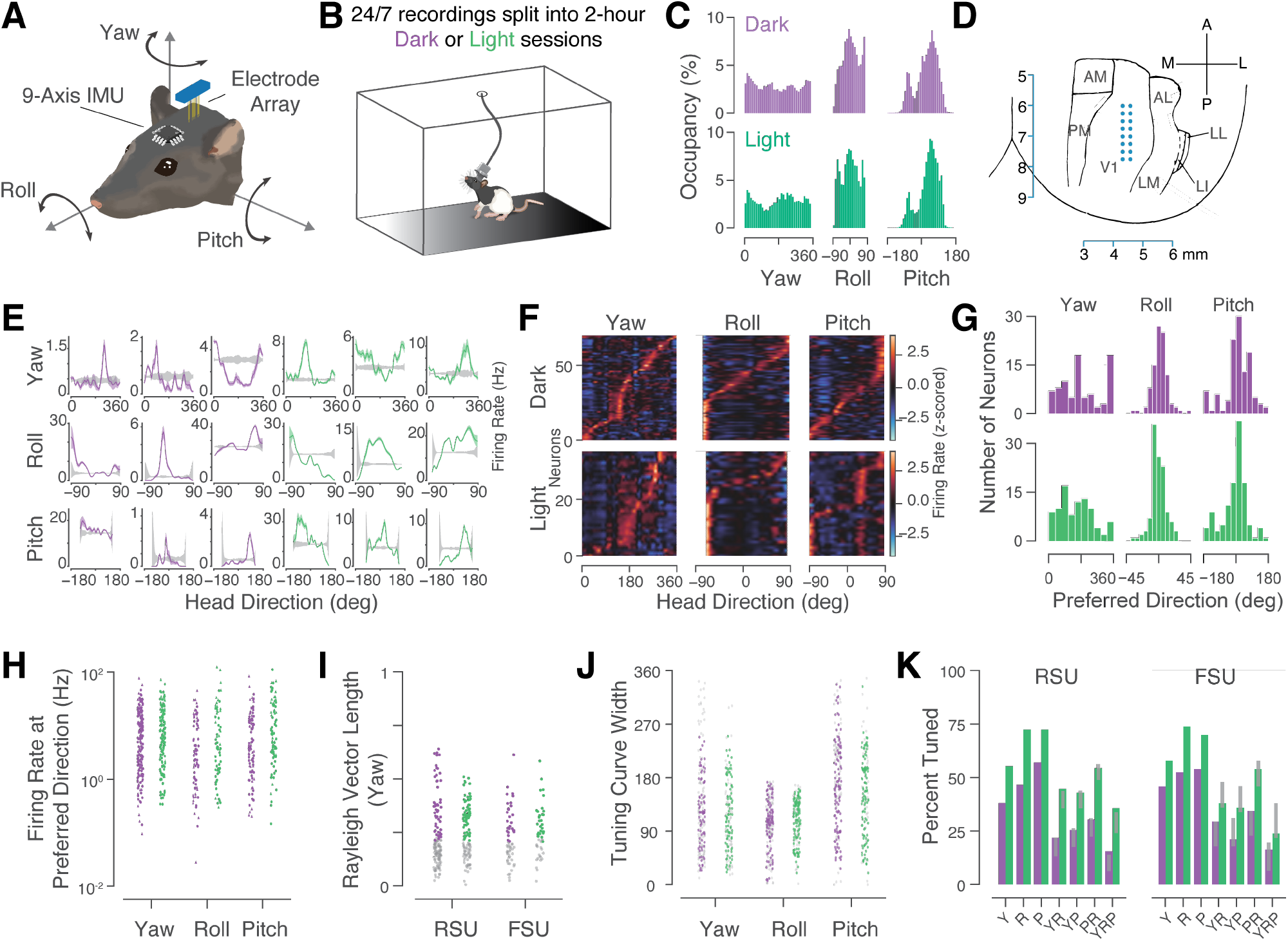
V1 Neurons are Tuned to 3D Head Direction in Freely Moving Rats. **(a)** Head direction (HD) angles were measured in 3D using a head-mounted sensor while neural activity in V1 was measured using chronically-implanted electrode arrays. **(b)** During the 24/7 recordings, rats were free to explore in their home cage either in the light or in the dark, with ∼12 hours per lighting epoch. Each epoch was then split into 2-hour sessions for analysis. **(c)** Behavioral coverage of the HD angles in dark (purple) and light (green) across rats and sessions. **(d)** Tetrodes were arrayed in 8×2 grids and implanted in V1 along the anterior-posterior axis. **(e)** Examples of tuning curves from 18 single units in V1 recorded in dark (purple) or light (green), with firing rates (mean ± SEM) plotted as a function of head direction along the yaw (top), roll (middle), and pitch (bottom) components. Grey shading: 95 CI of shuffle distributions. **(f)** 3D HD activity of 71 simultaneously-recorded V1 neurons in the dark (top) and 38 simultaneously recorded neurons in the light (bottom). Neurons were sorted by the angle corresponding to the highest firing rate for each given cell. **(g)** Preferred directions of all neurons with significant tuning. **(h)** Mean firing rates at preferred direction for all significantly tuned neurons were widely distributed in the dark across yaw (RSU: 9.67 ±8.7 Hz (mean ± std); FSU: 7.47 ±8.5 Hz), roll (RSU: 11.24 ± 10 Hz; FSU: 16.44 ±22 Hz), and pitch (RSU: 12.92 ±11 Hz; FSU: 19.69 ±21 Hz); and in the light across yaw (RSU: 10.98 ±13 Hz; FSU: 19.84 ±15 Hz), roll (RSU: 10.56 ±12 Hz; FSU: 23.64 ±16 Hz), and pitch (RSU: 16.11 ±20 Hz, FSU: 27.38 ± 20 Hz). Colored points represent significantly-tuned neurons; gray points: untuned neurons. **(i)** Distribution of Rayleigh vector lengths (RVLs) for neurons significantly tuned to yaw in the dark (RSU: 0.38 ± 0.017 (mean ± sem; FSU: 0.33 ± 0.017) and light (RSU: 0.34 ± 0.0099; FSU: 0.32 ± 0.017) Gray: non-significant neurons. **(j)** Distribution of tuning curve widths for neurons significantly tuned to yaw (RSU: 135.14 ± 7.51 degrees (mean ± sem); FSU: 97.53 ± 10.14), roll (RSU: 105.97 ± 5.29; Roll FSU: 98.65 ± 6.58), and pitch (RSU: 161.23 ±8.77; FSU: 162.48 ± 12.25) in the dark; and yaw (RSU: 118.96 ± 7.05; 122.07 ± 11.16), roll (103.35 ± 4.05; FSU: 116.41 ± 5.39), and pitch (RSU: 142.48 ± 7.02; FSU: 168.21 ± 10.95) in the light. **(k)** Percentage of neurons tuned to individual components of 3D HD (*Y, R, P*) and conjunctive tuning to multiple components (*Y* × *R, Y* × *P, P* × *R, Y* × *R* × *P*). 54 of 142 RSUs (38%) and 28 of 61 FSUs (45%) recorded in the dark were tuned to yaw; 66 of 142 RSUs (46%)and 32 of 61 FSUs (52%) were tuned to roll; 81 of 142 RSUs (57%) and 33 of 61 FSUs (54%) were tuned to pitch. In the light, 62 of 112 RSUs (55%) and 29 of 50 FSUs (57%) were tuned to yaw; 81 of 112 RSUs (72%) and 37 of 50 FSUs (74%) were tuned to roll; 81 of 112 RSUs (72%) and 35 of 50 FSUs (70%) were tuned to pitch. Rate of tuning to multiple components was within the 95% CI (gray bars) of a shuffle distribution that assumed tuning to multiple components to be an independent process (i.e. *P*(*Y* × *R* × *P*) = *P*(*Y*) × *P*(*R*) × *P*(*P*)).

Animals explored a uniform range of the yaw component of 3D HD and concentrated the roll and pitch components of 3D HD around zero and 50 degrees below the horizon, respectively, as previously reported (Figures 1c, S1c) ^33,34^. While the animals tended to move somewhat more in the dark than in the light (Figure S1b), there were no systematic differences in behavioral coverage between dark and light sessions (Figure 1c).

To define the cellular basis for a possible 3D HD signal in V1, we spike-sorted well-isolated units recorded in dark or light sessions (*N* = 365 neurons in *N* = 5 rats) (Fig S2a,b), which were then classified into regular-spiking putative excitatory units (RSUs, *N* = 142 in dark; *N* = 112 in light) and fast-spiking putative inhibitory units (FSUs, *N* = 61 in dark; *N* = 50 in light) based on waveform shape (Fig S2c,d). Recording locations were verified posthoc using electrolytic lesions of recording sites (Figures 1d, S2e) and by assessing responses to visual stimulation (Figure S2f-h) ^23^.

V1 neurons showed clear tuning to individual components of 3D HD (Figure 1e-g), with mean firing rates peaking at particular directions along any given HD axis for many cells. Neurons were deemed significantly tuned to yaw if the directionality of their firing rate vector (Rayleigh vector length^35–38^) exceeded the 95% CI of a shuffle distribution (see Materials and Methods). For roll and pitch, neurons were classified as significantly tuned if the peak mean firing rate exceeded the 95% CI of a shuffle distribution. By these criteria, ∼40% were tuned to yaw in the dark and ∼56% in the light; ∼48% were tuned to roll in the dark and ∼72% in the light; and ∼56% were tuned to pitch in the dark and ∼71% in the light.

Neurons recorded simultaneously showed a wide degree of directional preference, spanning the extents of each plane (Figure 1f). Across all recorded cells that showed directional tuning, preferred directions were widely distributed for yaw, but concentrated for roll and pitch (Figure 1g). Of the neurons significantly tuned to pitch, ∼ 1/3 preferred negative values of pitch (i.e. head pointing up). Only 30% of cells had preferred pitch within 15 degrees of zero in the dark and 16% in the light; thus, the majority of neurons encoded angles that deviated from zero-degree pitch, suggesting that these neurons do not simply fire during any behavior confounded with zero-degree pitch. In contrast, preference for roll was tightly distributed, with ∼ 86% of cells tuned within ± 15 degrees from zero.

Mean firing rates at preferred directions spanned a wide range, similar to previous reports of cortical firing rates ^39^ and firing rates of HD cells in various brain regions ^14,30,32^ (Figure 1h). Rayleigh vector lengths (RVLs) for yaw were lower than in classical HD cells previously reported, but significantly above chance, ranging 0.21 − 0.64 compared to a shuffle distribution’s 95% upper CI of ∼ 0.2 (Figure 1i). RVLs were somewhat higher for RSUs (0.36 ± 0.01 (mean ± sem)) than FSUs (0.32 ± 0.01) (*p* = 0.016, Mann-Whitney U (MWU) test), but not significantly different in the dark (0.36 ± 0.01) versus light (0.33 ± 0.01) (*p* = 0.13 MWU test). Tuning curve widths were similar to previous reports of HDs in the rat lateral mammillary nuclei (LMN)^40^ and dorsal presubiculum of Egyptian fruit bats ^35^, but wider than the rat postsubiculum^41^ (Figure 1j).

While ∼ 50% of all neurons were tuned to individual components of 3D HD, we next asked whether individual neurons represented 3D HD, or whether 3D tuning arose on a population level. To assess the possibility of conjunctive tuning, we calculated the proportion of neurons tuned to multiple components of 3D HD (Figure 1k). The proportion of neurons tuned to two components (e.g. yaw and roll or roll and pitch) was ∼ 25%, while the proportion tuned to all three was ∼ 10% in the dark and ∼ 25% in the light. All of these fell within the 95% CI of a shuffle distribution in which the identity of the tuned neurons was randomly permuted 1000 times. Thus, the proportion of neurons tuned to multiple components of 3D HD was similar to what could be expected if tuning to components were independent, indicating that even though V1 represents 3D HD as a population, individual V1 neurons likely receive information about the components of 3D HD from independent sources.

### V1 Multiunit Ensembles Encode 3D Head Direction

To address whether V1 population dynamics reflect 3D HD on a mesoscopic scale, we built linear regression models that took as inputs the multiunit activity (MUA) firing rates from the 16 tetrodes, and predicted the HD in the three Euler angles separately (Figures. 2a-c, S3a-c) (see Materials and Methods). We reasoned that MUA firing rates, which were calculated by binning times of all spikes extracted from each tetrode, could serve as a proxy for coordinated V1 population activity^42^. The models performed well above chance, both in the dark and in the light (Figure 2d,e). Decoding performance was highest for the roll and pitch components and did not differ significantly between the light and dark conditions for any of the planes (Figure 2d,e). Decoding performance increased as a function of MUA window size, and was generally highest when the lag between neuronal activity and HD was close to zero milliseconds (Figure 2f,g).

**Figure 2:**
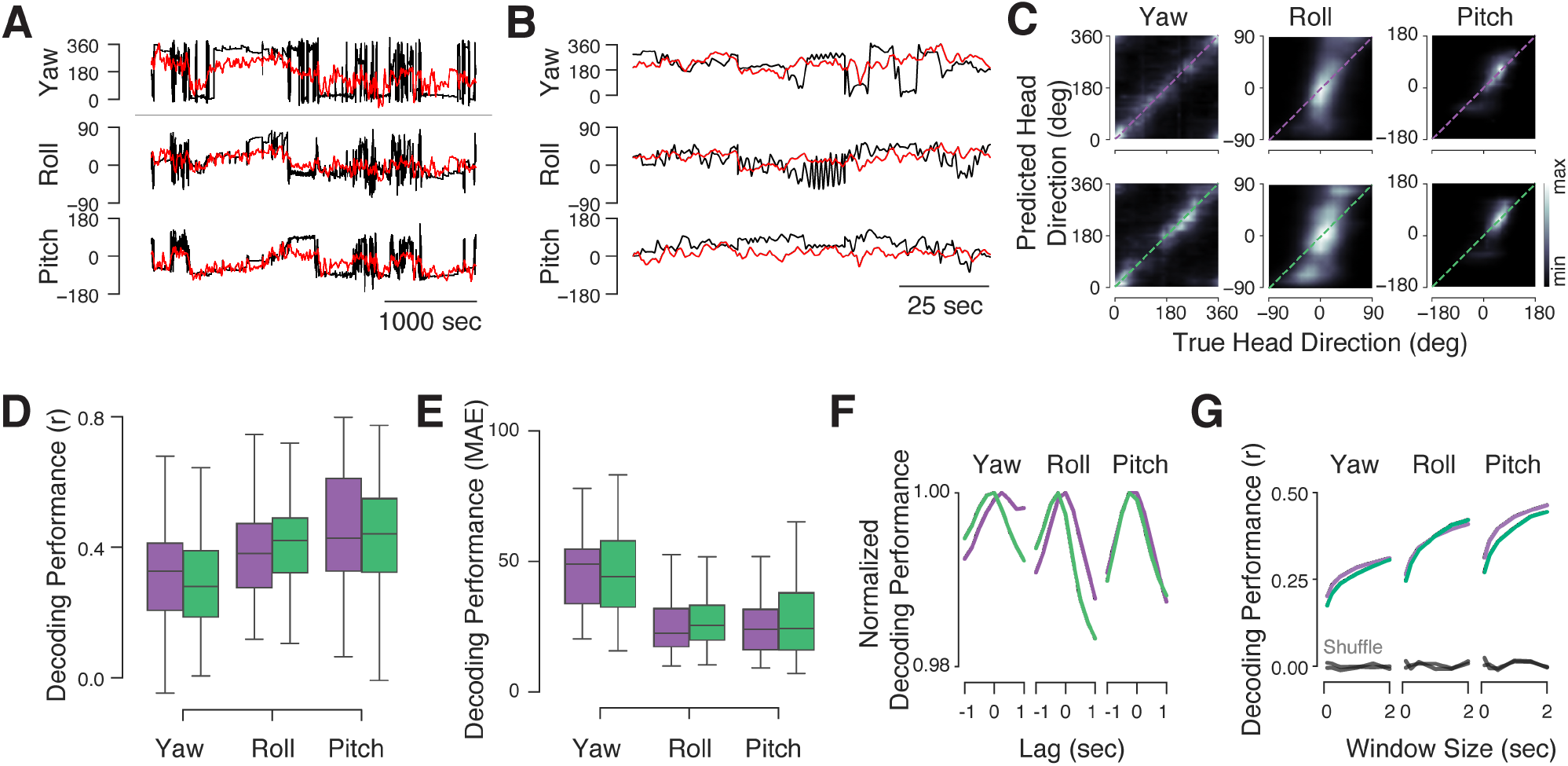
V1 Multiunit Ensembles Encode 3D Head Direction. **(a**,**b)** Example traces from one decoding session showing true (black) and predicted (red) HD angles in the dark. Decoding was performed using linear ridge regression with multiunit activity (MUA) firing rates as inputs. **(c)** Mean bivariate histograms of true and predicted HD angles across rats and sessions in the dark (top row) and light (bottom row). Diagonal lines indicate ideal decoding performance. **(d)** Summary of decoding model performance measured using the circular correlation coefficient for yaw models and Pearson’s correlation coefficient for the roll and pitch models. Green: light; purple: dark. Yaw, dark: *r*_*circ*_ = 0.31 ± 0.0154, light *r*_*circ*_ = 0.30 ± 0.0154 (mean ± SEM); Roll, dark *r* = 0.41 ± 0.0162, light *r* = 0.42 ± 0.0144; Pitch, dark *r* = 0.46 ± 0.0182, light *r* = 0.44 ± 0.0167 (mean ± s.e.m.). **(e)** Performance of same models measured using median absolute error (MAE). Yaw, dark: MAE=45.53 ± 1.5038 vs. shuffle (not shown) MAE=69.18±1.6823 (mean ± sem), *p* = 3.4*e*−17 MWU test; light: MAE=46.24 ±1.7557 vs. shuffle MAE=64.37 ± 2.1038, *p* = 1.3*e* − 09 MWU test. Roll, dark: MAE=26.60 ± 1.4475 vs. shuffle MAE=32.84 ± 1.4954, *p* = 0.0004 MWU test; light: MAE=27.80 ± 1.1347 vs. shuffle MAE=33.68 ± 1.2500, *p* = 0.00014 MWU test. Pitch, dark: MAE=24.93 ± 1.0577 vs. shuffle MAE=35.21 ± 1.7982, *p* = 5*e* − 05 MWU test; light: MAE=26.61 ± 1.3740 vs. shuffle MAE=34.45 ± 1.9417, *p* = 0.0031 MWU test. **(f)** Decoding performance as a function of MUA window size for dark or light sessions, or those in which the HD traces were randomly permuted before model fitting (grey). Window lag was 0 seconds for these models. **(g)** Decoding performance as a function of window lag while window size was held constant at 500 ms. Negative lags represent cases where neural activity occurs before a given HD point; positive lags indicate cases where neural activity occurs after a given HD point.

In previous work, we reported that V1 responses to head orienting movements (HOMs) depend on secondary motor cortex (M2) ^23^. To address the possibility that V1 HD encoding also depend on M2, we fit and tested the linear regression models to sessions recorded in M2-lesioned rats. These models performed well in the dark and the light, for yaw, roll, and pitch, albeit with reduced performance for pitch relative to non-lesioned animals (Figure S3d). Thus, neuronal dynamics in V1 encode the three angular components of HD in freely behaving rats, in a manner that likely does not depend on inputs from M2.

### Decoding 3D Head Direction from V1 Single Unit Activity

While V1 multiunit populations encoded the 3D HD signal and single units were tuned to components of 3D HD, it is unclear if populations of individual V1 neurons could encode 3D. To assess this possibility, we constructed classification models that predicted binned direction of individual components of 3D HD using firing rates from single unit activity concatenated into pseudopopulations ^43^.

These models performed with high accuracy (Figure 3a). Classification was highest for models fit and tested using both RSUs and FSUs, and was higher for RSUs than FSUs, suggesting that V1 pyramidal neurons have better access to directional information than local interneurons (Figure 3b). Models fit on small fractions of neurons performed close to chance, and performance steadily increased as more neurons were included (Figure 3c). Model performance likewise improved with increasing size of the firing rate temporal window, indicative of a possibility that V1 neurons might integrate HD information on the timescale of seconds (Figure 3d).

**Figure 3:**
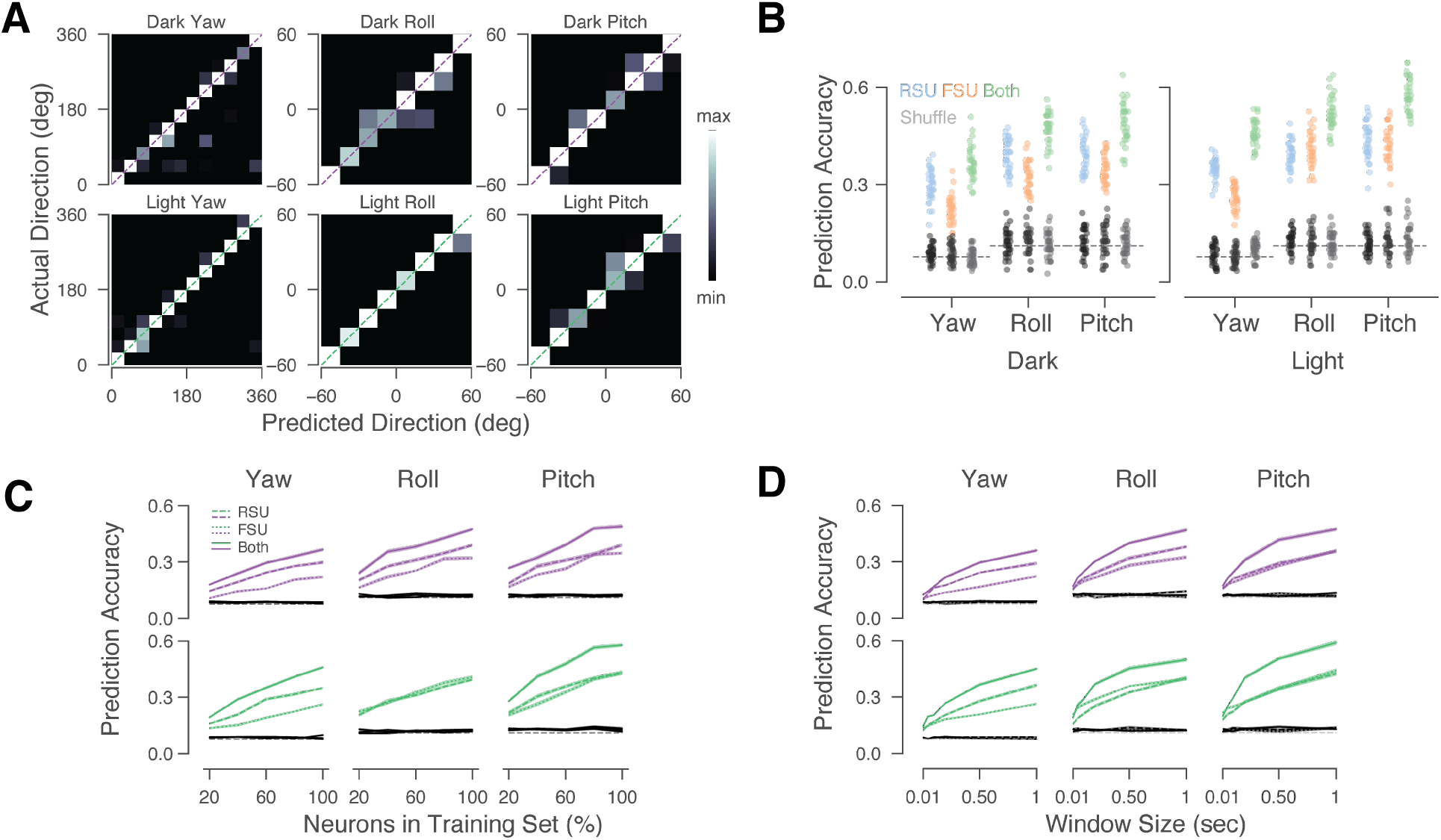
Populations of V1 Single Units Encode 3D Head Direction. **(a)** Confusion matrices from logistic regression models fit and tested on single-unit data. The yaw signal was binned in 30-degree bins; the roll and pitch were binned in 15-degree bins. The models were fit with 90% of the samples and tested on the remaining (unseen) 10%. **(b)** Decoding accuracy (number of correctly-predicted labels out of all labels in testing set) for models fit with data from all neurons, with a temporal window of 1 second, split by neuron type (RSU: blue; FSU: orange; both types combined: green). Gray: models fit with shuffled data. Dotted lines: expected chance performance (1 / number of categories). **(c)** Decoding accuracy as a function of the number of neurons used to fit and test the models, split by the neuron type (RSU: dashed; FSU: dotted; both types combined: solid lines). **(d)** Decoding accuracy as a function of the temporal window size, split by neuron type as in **c**.

### Head Direction Tuning is Modulated by Overall Movement

Previous work has demonstrated that movement has a profound effect on sensory cortical dynamics, even in the absence of sensory stimuli ^18,22,23,44^. To examine if V1 HD signals are modulated by the animal’s movement, we split the recordings into moving and resting chunks based on the profile of the total acceleration of the head (Figure 4a) (see Materials and Methods).

**Figure 4:**
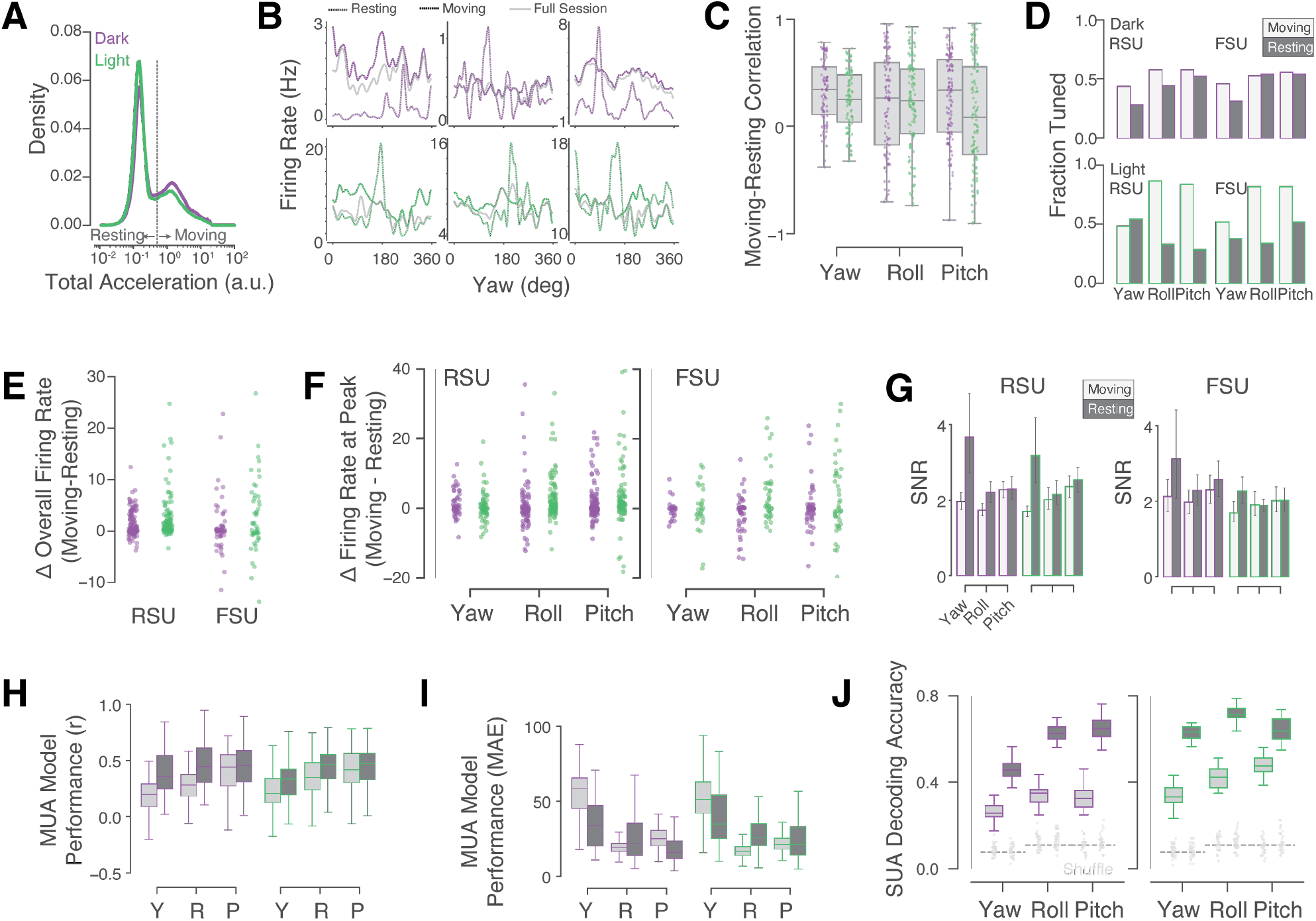
Head Direction Tuning is Modulated by Overall Movement. **(a)** Total acceleration (the norm of linear acceleration along the three planes) was used as a proxy for the animal’s overall movement. Sessions were split into moving or resting chunks based on the total acceleration, which showed a bi-modal distribution. **(b)** Example yaw tuning curves for six neurons in the dark (top row) and light (bottom row) split by movement condition. **(c)** Tuning curves calculated during movement or rest were only mildly correlated (r=0.22 ±0.01, mean ± sem)). **(d)** A moderately higher fraction of neurons were tuned to components of 3D HD during movement (light gray) compared to rest (dark gray). **(e)** Difference between mean firing rates during movement vs rest for individual significantly-tuned neurons was skewed toward higher firing rates during movement. **(f)** Difference between firing rates at preferred direction during movement compared to rest for individual significantly-tuned neurons. **(g)** The signal-to-noise ratio (SNR) during movement (light gray) vs rest (dark gray), calculated as the mean firing rate at preferred direction divided by overall firing rate across all directions for individual neurons. SNRs were higher during rest than movement. **(h**,**i)** Ridge regression MUA decoding model performance as Pearson’s r (**h**) and the mean absolute error (MAE) between predicted and actual HD (**i**) for models fit and tested on moving or resting data. **(j)** Accuracy of logistic regression SUA decoding models fit and tested on moving or resting data.

Animals spent 40% or 60% of time in the moving or resting state, respectively. Nevertheless, the animals covered the distribution of head orientations at comparable rates in the two conditions (Figure S4a). The distributions of moving or resting bout durations were wide, with prominent peaks around the median duration of ∼1 second (Figure S4b). Overall, the animals spent ∼3% of the time resting for one minute or longer (Figure S4c). In contrast, previous studies of the distribution of sleep bouts in rats reported that sleep bouts last ∼16-20 minutes ^45–47^. Thus, while the resting condition necessarily included bouts of sleep in addition to quiet wakefulness, the small fraction of time the animals spent resting at durations compatible with sleep allowed us to directly compare tuning curves in the moving and resting conditions.

Head direction tuning curves of individual cells tended to be different during movement compared to rest, with moderate correlations between the two states (Figure 4b,c). Slightly smaller fractions of V1 neurons were tuned to components of 3D HD during rest compared to movement (Figure 4d). However, both the mean and peak firing rates were higher during movement than rest for both RSUs and FSUs (Figure 4e-f). Overall, the tuning curve signal-to-noise ratios (SNRs)–defined as the firing rate at the preferred direction over the mean background firing rate–was slightly higher during rest than movement (Figure 4g), likely due to the increased overall firing rate during movement. Correspondingly, decoding models performed better during rest than movement for both the MUA data (Figure 4h,i) and SUA data (Figure 4j). Thus, movement eroded encoding of 3D HD in V1, likely via an increase in background firing.

### Head Direction Tuning in V1 is Stable

To be useful for navigation, it would be beneficial for HD representations to be stable over time and invariant to environmental conditions. V1 neurons significantly tuned to HD exhibited stable tuning curves and low discrepancy in preferred directions in the two halves of each recording session (Figure 5a-c). Further, the discrepancy in preferred direction was related to the RVL, with stronger tuning corresponding to lower discrepancies in the two halves of a session (Figure 5d). Thus, V1 neurons exhibited basic properties of HD tuning stability.

**Figure 5:**
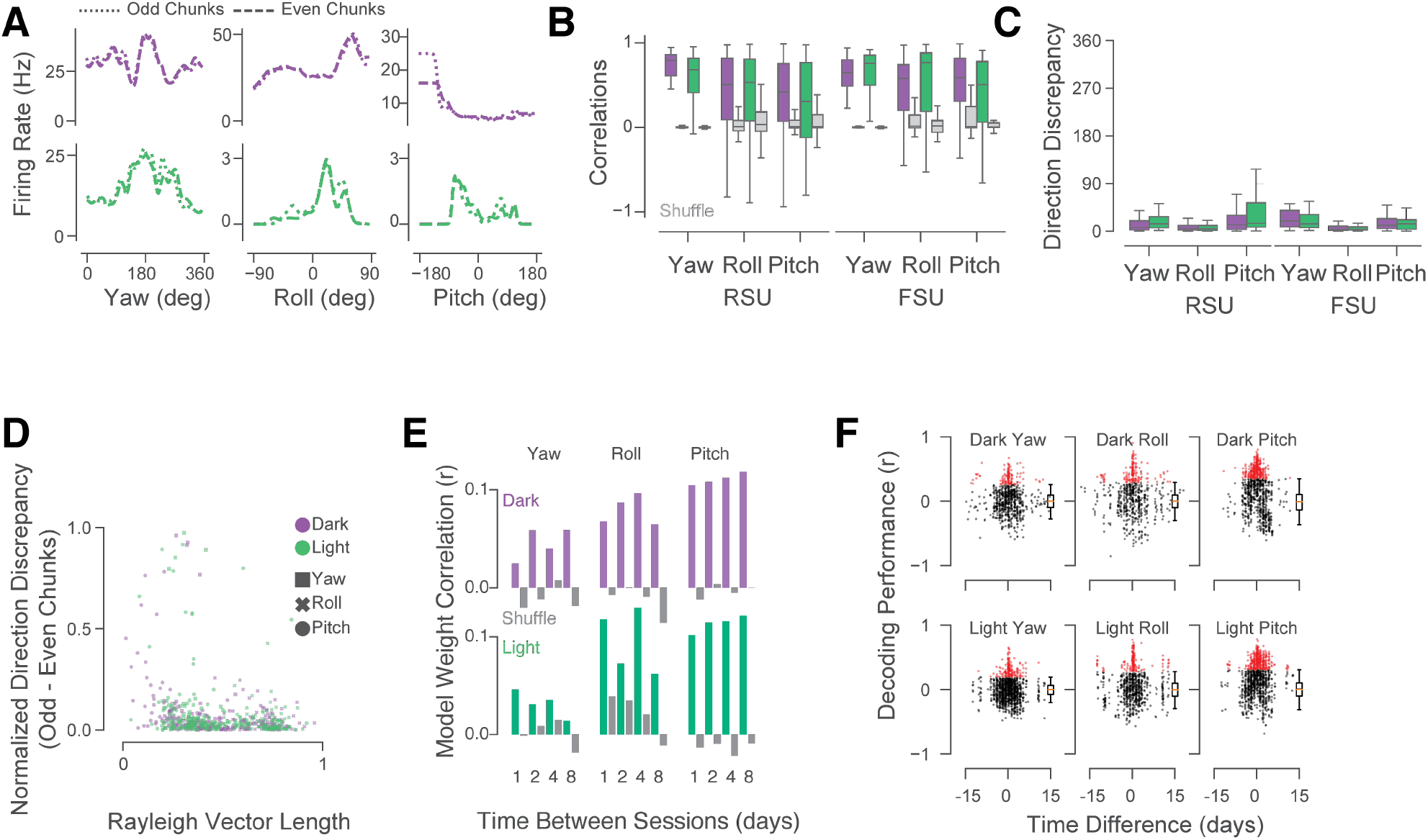
Encoding of Head Direction by V1 Neurons is Stable. **(a)** Example head direction tuning curves splitting the data from single sessions into halves (30-second non-overlapping odd and even chunks). **(b)** Correlations between odd and even chunks of neurons tuned to components of 3D HD. **(c)** Differences in preferred direction between odd and even chunks of single sessions of neurons significantly tuned to HD. **(d)** Normalized differences in preferred direction as a function of the Rayleigh vector length. Neurons with stronger tuning (longer vector length) tended to have smaller discrepancies in preferred direction between odd and even session chunks (Spearman’s *r* = −0.30, *p* = 4.6 × 10^−5^). **(e)** Correlations of MUA ridge regression model weights among session pairs separated by 0-24, 24-48, 48-96, or 96-192 hours. Purple (dark) and green (light) indicate session pairs for one given rat (i.e. within-rat comparisons); grey bars indicate correlations among session pairs from different rats (i.e. across-rat comparisons). Sessions with above-median decoding performance were used for the within-rat comparison. **(f)** Decoding performance of models trained on MUA data from one session and tested on other sessions from the same animal, plotted as a function of time between the fit and test sessions. Each dot represents the performance (correlation coefficient) of one fit-test pair; boxplots show distribution of models tested on firing rates from other rats (control); red dots indicate sessions with decoding performance above the 95% CI of the control distributions.

To address how visual inputs may impact HD tuning of individual neurons in V1, we tracked single units between the dark and light conditions for a subset of experiments (Figure S5a-c). Mean firing rates were different between the dark and light conditions ^23^ and tuning curves of many neurons were moderately similar across the two lighting conditions (Figure S5d,e). While ∼ 25-50% of neurons in these recordings were tuned to HD in either dark or light, ∼ 12-24% were tuned to HD in both conditions; these rates were comparable to a null distribution in which cell identity was shuffled, suggesting that V1 neurons were tuned to HD independently in the two lighting conditions (Figure S5f). Examining the differences in preferred direction of neurons tuned to HD in both light and dark revealed that most neurons had low directional discrepancy between light and dark, although the drift was higher for yaw, as has previously been reported for recordings performed in the dark in subcortical head direction circuits^14,48^ (Figure S5g).

To examine the longterm stability of the HD signal in V1, we analyzed the correlation structure of weights from the MUA linear regression models used to decode 3D HD over time. Each session’s weights (200 time-points per tetrode × 16 tetrodes) were correlated to weights from sessions from the same rat separated by 0-1, 1-2, 2-4, or 4-8 days, and compared to correlations of weights taken from different rats (Figure 5e). We reasoned that weight correlations from across rats should serve as a control measure, since there is no reason to expect tetrodes from different brains to be weighed similarly. Within-rat weight correlations were higher than across-rat correlations for all models, and especially for the roll and pitch, in both dark and light.

To more directly address the stability of 3D HD encoding in V1, we next performed cross-session decoding, in which models fit on data from one session were tested on data from the same rat’s other sessions, separated by up to 15 days (Figure 5f). Our calibration tests on the IMU showed that it was capable of stable measurements of the roll and pitch angles, but tended to drift for the yaw (Figure S1d-f), suggesting that yaw angles would not accurately transfer between sessions; nevertheless, decoding models could be used across sessions because the performance measure for yaw, the circular correlation coefficient, is robust to such offsets or drift as long as all values drifted coherently. While cross-session decoding performance was highly variable and the highest performance was achieved from models fit closest in time to the test session, many sessions could be decoded above chance up to a week later, indicating that the structure of the underlying signal was not systematically drifting (Figure 5f). Together, these results indicate that while the directional tuning of individual V1 neurons may drift somewhat in the absence of visual cues, the multiunit signal that formed the basis for the linear decoding of 3D HD was stable on the order of a week.

## Discussion

Using chronic electrophysiological recordings in freely behaving rats, we showed that primary visual cortex encodes three-dimensional components of head direction, or the orientation of the head in 3D space, either in the presence or absence of visual cues. This signal was modulated by movement and lighting conditions, but on a mesoscopic level remained stable across time.

While several previous studies have failed to find HD signals that extend beyond yaw angle (azimuth) in the rodent limbic system^37,49^, recent work has validated the idea that navigational variables represent three-dimensional space^50^. 3D HD cells were found in the presubiculum of crawling or flying bats ^35^, and 3D place cells were found in both bat and rat hippocampus^51,52^. It is not entirely clear why some studies have failed to find 3D HD cells in the rodent limbic system, but one possible explanation is methodological: rats in the present study were free to move as they wished in the 2D arena while the head was free to point anywhere in the 3D space, while in some previous studies animals were required to crawl upside down along a ceiling^14^ or vertical walls^49,53–55^ or restrained and moved passively^37^. While V1 encoded 3D HD as a population, the neurons analyzed here were tuned to multiple components of 3D HD at rates expected by chance if the components were independent, suggesting that yaw, roll, and pitch signals may arrive in V1 from independent sources. Further work into synaptic tracing of HD signals and naturalistic behavior in 3D space may resolve these issues.

As far as it is possible that some neural responses were correlated with particular behaviors in the cage, only drinking from the water bottle would have occurred in a narrow range of orientations. The water bottle was the only item in the cage fixed in place; the toys and food bowl were free for the rats to move around, which they did frequently. While we cannot definitively rule out the possibility that some neurons may have responded to drinking, drinking alone would not explain the range of directions to which the neurons were tuned to in yaw, roll, and pitch. While our data do not allow us to disambiguate the rat’s yaw and position, we believe the observed yaw tuning is not affected by position because the data come from long recording sessions and are averaged over two-hour analysis chunks; it would be improbable that a rat’s yaw and position were correlated on such timescales. Furthermore, tuning to yaw was stable for odd and even 30-second chunks of the two-hour analysis chunks, suggesting that immobility at particular corners of the cage was unlikely to influence yaw tuning. Finally, previous reports have shown that rats sleep in bouts of 16–20 min^45–47^. Thus, while it is possible that sleep activity and yaw were somewhat correlated in our analysis, this could not explain the variety of represented yaw directions by simultaneously recorded neurons (Figure 1f). Our observation of tuning to yaw during movement further contributes to the idea that yaw tuning was not a confound of sleep activity.

Preferred angles for roll and pitch centered around zero degrees. One potential explanation for such concentrated tuning around the orientations at which the animals spent the most time (see Figure 1c) is that cortex may be maximizing the efficiency of information encoding^56–58^. Roll and pitch are egocentric measures, compared to the yaw/azimuth, which is allocentric. It is possible that roll and pitch reflect the animal’s sense of posture rather than the direction its head is pointed in per se, and coding for the latter has previously been reported in other brain areas ^40,59,60^. Our data cannot disambiguate these two possibilities and it is possible, of course, that the tuning curves for roll are simply a product of neurons being suppressed during behaviors corresponding to negative or positive rolls of the head; in either case, the firing patterns of these neurons accurately represent the roll.

While single units exhibited mild stability between dark and light sessions (Figure 5a-c), our data cannot address the stability of individual V1 neurons on the timescale of days or weeks. Others have previously reported drift on the order of 25-30 degrees in tuning to yaw HD during short (∼8 minute) recording sessions ^48,61^. Our data show larger drift (∼90 degrees for yaw; ∼10 degrees for roll; ∼50 degrees for pitch; see Figure S5g) over the course of two-hour sessions. The discrepancy may be due to the longer recording sessions or drift in the IMU’s yaw readings; alternatively, competing inputs to the visual system may also play a role in shaping tuning to HD. Nevertheless, on a mesoscopic multiunit level, populations of V1 neurons were capable of encoding HD in a stable manner over the course of a week (Figure 5e,f).

While the head direction information that we have decoded could reasonably be the product of visual processing in the presence of light, the fact that these signals are also present (and stable) in the dark points to an exogenous source of this information. Retrosplenial cortex (RSC) is one plausible candidate for this source, as it processes vestibular information and sends extensive projections to V1^19^. In previous work, we demonstrated that rat V1 responds to orienting movements of the head in a direction- and light-dependent manner ^23^. We found also that V1 responses to such movements were severely reduced following lesions to secondary motor cortex (M2), even though the movements themselves were relatively unaffected. Applying the HD decoding models to M2-lesioned animals showed that V1 in these animals still encoded 3D HD (Figure S3d), suggesting that HD signals in V1 originate in a different region.

Our study is technically not the first to find HD representation in the visual system. Chen et al. (1994) found HD cells in a higher visual cortical area, Oc2M^30^. HD cells in the visual cortex have also recently been recorded using a head-mounted two-photon calcium imaging microscope^62^. Notably, while the observed RVL values in the present study were lower than reported in classic HD brain regions, they were on par with those from the two-photon visual cortex data^62^.

What purpose, if any, might head direction representation in a primary visual area serve? One possibility is that head-direction representations in visual cortex play a role in reconciling visually-driven and vestibular contributions to an overall representation of head direction. Elegant theoretical work has posited a ring attractor as a possible computational substrate for estimating head direction in the brain, with input from multiple modalities ^63^; a literal ring of neurons implementing a ring attractor that computes head direction has recently been described in the brains of fruit flies ^3–6^. While our results do not definitively speak to the presence of such a system in the brain of the rat, the presence of stable, persistent representations of heading from multiple modalities could play a role in such a system if it exists.

Another, non-mutually-exclusive, possible explanation for HD signals in V1 is that these are used by visual circuits to aid in predicting future visual features. This paradigm proposes that a core function of the visual cortical hierarchy is to make predictions about future states ^29,64–66^. The finding here that primary visual cortex contains information about head direction does not mean that these cells are necessary for navigation per se; it does however suggest an intriguing possibility that navigational signals like HD may be used by the visual cortex to improve predictions about future visual features, as the animal moves through the environment. Future work may test this idea experimentally *in vivo*, for example by isolating and manipulating V1-projecting axons containing HD information in the context of a behavioral task.

While it is broadly accepted that vision plays a significant role in shaping the brain’s sense of space and heading direction, little work has been done to ask how the computation of such cognitive variables impacts dynamics in sensory areas. We have taken a step toward elucidating the interplay of visual sensory signals and those that encode head direction. Future work will investigate exactly where these signals originate, and how they impact sensory perception and behavior. The present work adds to a growing awareness that non-visual signals play a critical role in shaping visual cortical population dynamics, and that the computational role of visual cortex may extend beyond just processing visual inputs. Understanding how the visual cortex interacts with multimodal systems of the brain, such as those used in navigation and spatial orientation, holds the promise to deepen our understanding of cortical computation in general.

## Materials and Methods

### Animals

The care and experimental manipulation of all animals were reviewed and approved by the Harvard Institutional Animal Care and Use Committee. Experimental subjects were female Long Evans rats 3 months or older, weighing 300-500 g (*N* = 9, Charles River, Strain Code: 006).

### Surgery

Rats were implanted with 16-tetrode electrode arrays targeting L2/3 of V1 in the right hemisphere, as described previously^23^. Animals were anesthetized with 2% isoflurane and placed into a stereotaxic apparatus (Knopf Instruments). Care was taken to clean the scalp with Povidone-iodine swabsticks (Professional Disposables International, #S41125) and isopropyl alcohol (Dynarex #1204) before removing the scalp and cleaning the skull surface with hydrogen peroxide (Swan) and a mixture of citric acid (10%) and ferric chloride (3%) (Parkell #S393). Three to four skull screws (Fine Science Tools, #19010-00) were screwed into the skull to anchor the implant. A 0.003” stainless steel (A-M Systems, #794700) ground wire was inserted ∼ 2mm tangential to the brain over the cerebellum.

Tetrodes were arrayed in 8×2 grids with ∼ 250-micron spacing, and were implanted in V1 with the long axis spread along the AP (ranging 6-8 mm posterior to bregma, 4.5 mm ML, targeting layer 2/3 at 0.6 mm DV). The dura was glued (Loctite) to the edges of the craniotomy to minimize movement of the brain relative to the electrodes. After electrodes were inserted into the brain, the craniotomy was sealed with Puralube vet ointment (Dechra) and the electrodes were glued down with Metabond (Parkell). Post-operative care included twice-daily injections of buprenex (0.05mg/kg Intraperitoneal (IP)) and dexamethasone (0.5 mg/kg IP) for three days.

### Behavior

Spontaneous behavior in rats living in a 15×24” home cage was recorded under two conditions: dark, in which the lights in the box and room were turned off, and light, in which the box was illuminated. Recordings were carried out 24/7 and split into 2-hour sessions in dark or light on a pseudo-random light cycle, as previously described^23^.

The behavior box was constructed from aluminum extrusions and black extruded acrylic (McMaster). The floor was covered in bedding and the arena contained a cup with food, a water bottle and toys. The walls were lined with strips of white tape at different orientations to provide visual features in the light condition, and the box was outfitted with white LED strips (Triangle Bulbs Cool White LED Waterproof Flexible Strip Light, T93007-1, Amazon) to provide illumination.

For recordings, rats were tethered with a custom 24” cable (Samtec, SFSD-07-30C-H-12.00-DR-NDS, TFM-107-02-L-D-WT;McMaster extension spring 9640K123) to a commutator (Logisaf 22mm 300Rpm 24 Circuits Capsule Slip Ring 2A 240V TestEquipment, Amazon). A 9-axis Inertial Measurement Unit (IMU) (BNO055, Adafruit) was used to record movement; the sensor was epoxied to the connector on the cable, in a way that placed it directly above the electrodes and headstage. This not only ensured that the sensor was always in the same position above the animals’ heads, but also that it stayed powered after the animals were unplugged, preventing the need to re-calibrate the sensor after each recording. The IMU data were acquired at 100Hz using a micro-controller (Arduino) and saved directly to the acquisition computer’s disk. To synchronize IMU and electrophysiology data, the Arduino provided a 2-bit pseudo-random pulse code to the TTL inputs on the electrophysiology system.

### Electrophysiology

Tetrodes were fabricated using 12.5-micron nichrome wire (Sandvik-Kanthal) following standard procedures ^67–69^, as described previously^23^. Tetrodes were threaded through 42 AWG polyimide guide tubes into 8×2 grids of 34 AWG tubes (Small Parts) and glued to a single-screw micro-drive. The drive was modified from a design in Mendoza et. al^70^ and Vandercasteele et. al^71^, in which a 3-pin 0.1” header served as the skeleton of the drive, with a #0-80 screw replacing the middle pin, and the header’s plastic serving as the shuttle. In the experiments reported here, the tetrodes were not advanced after recording sessions started. The tetrodes were plated with a mixture of gold (Neuralynx) and polyethylene glycol (PEG) as per Ferguson et. al^72^, to an impedance of ∼100 − 250*K*Ω. The ground and reference wires were bridged and implanted through a craniotomy above the cerebellum.

Electrode signals were acquired at 30 kHz using custom-made Intan-based 64-channel headstages ^73^ and Opal-Kelly FPGAs (XEM6010 with Xilinx Spartan-6 ICs). Spikes were extracted following procedures described in Dhawale et al (2017)^73^. Multiunit firing rates were estimated in non-overlapping 10-ms bins from extracted spikes. Multiunit and single-unit firing rates were Gaussian-filtered and in some cases z-scored. Single-units were sorted using MountainSort ^74^ and classified into putative excitatory regular-spiking units (RSUs) or putative inhibitory fast-spiking units (FSUs) based on the trough-to-peak time (width) and full-width at half-max (FWHM) of the unfiltered waveforms (Figure S3).

We isolated *N* = 482 single units in *N* = 5 rats extracted from 24/7 recordings in chunks of two-hour sessions. Multiunit V1 recordings were also made from an additional *N* = 4 rats with M2 lesions. The spike-sorted two-hour-sessions were recorded in the dark (*N* = 10), light (*N* = 10), or during flashing (*N* = 5), where the cage lights were repeatedly flashed at random intervals to assess the responsiveness of the recorded neurons to visual stimulation (Figure S2). In the dark, we isolated *N* = 203 units (*N* = 142 RSUs, *N* = 61 FSUs). In the light, we isolated *N* = 162 units (*N* = 112 RSUs, *N* = 50 FSUs). In “flash” sessions used for testing visual responsiveness, we isolated *N* = 112 units (*N* = 78 RSUs, *N* = 34 FSUs). In a separate analysis, we simultaneously spike-sorted units (*N* = 100 neurons from *N* = 8 sessions) that were recorded in the dark or light, in order to track the influence of visual cues on encoding of HD in V1.

### Head Direction Tuning Curve Analysis and Significance Testing

Tuning curves were constructed by plotting mean activity in 3° bins, ranging 0-360° for yaw, −90-90° for roll, and −180-180° for pitch. The resulting tuning curve was then smoothed with a Gaussian filter with a standard deviation of 6° (2 bins). Preferred direction was calculated by taking the mean of the firing rates multiplied by the angles in the complex plane, and converting the resulting angle back to degrees. Rayleigh vector length (RVL) was used to calculate the directionality of each neuron’s firing properties in yaw and was defined as follows ^35^:

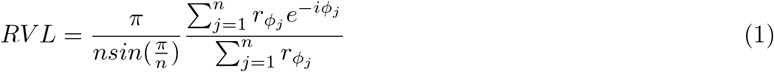

Here, *N* was defined as the number of head direction bins; 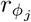 as the mean firing rate at head direction *ϕ*_*j*_; and *j* as the index of the binned head direction and mean firing rate vectors. The firing rates going into the RVL calculation were z-scored and minimum-subtracted in order to remove variation resulting from the vastly different background and peak firing rates (see Figure 1h) showing the distribution of firing rates at preferred directions). Significance testing was performed by comparing the RVL each neuron to a shuffle distribution that was calculated by randomly permuting the sequence of firing rates relative to the head direction vector 1000 times, calculating the shuffle tuning curve and taking the RVL of each shuffled tuning curve. A neuron was designated as significantly tuned to yaw if its RVL exceeded the 95% CI of the shuffled RVLs (a value ∼ 0.2, as previously reported^35,75^).

Neurons were classified as significantly tuned to the roll or pitch of HD if the mean firing rate at the preferred direction exceeded the 95% CI of the shuffled distribution^60^.

Tuning curve width was calculated by finding direction bins on either side of the preferred direction that were closest to the quarter of the height of the curve. The height of the curve was defined as the difference between the firing rate at the preferred direction and the minimum firing rate.

### Head Direction Decoding

Head direction in 3D was decoded separately for the yaw, roll, and pitch angles using Ridge linear regression for MUA data, and Logistic Regression for the SUA data. Because the yaw signal is circular, it was modeled in the complex plane (i.e. angles were decomposed into x and y components, fitted to two separate models, and recombined into angles during evaluation).

The Ridge models took z-scored MUA firing rates as inputs and predicted the angle of a given component of 3D HD. The MUA firing rates from *N* = 16 tetrodes were taken in time windows and flattened before fitting, such that each window x tetrode became a vector of features used to predict a HD angle (Figure S2).

The models were fit using half of the data in a session, with the session split into 30-second chunks separated by 0.25-second discarded gaps. The even chunks were used for fitting the linear regression models and the odd ones were used for testing. The Ridge models were implemented using the Scikit Learn Python package^76^, and cross-validated threefold. Each two-hour session was modeled separately (*N* = 89 sessions in dark, *N* = 95 sessions in light across *N* = 5 rats).

To determine optimal window size and lag, we performed a grid search, with window size ranging from 0.10, 0.25, 0.50, 1.00, 1.50, to 2.00 seconds, and window lag ranging from −1.00 to 1.00 seconds in steps of 250 ms. Negative lags represent neural activity lagging behind a given HD position (Figure 1i,j).

The logistic regression models were used to classify binned HD data based on population SUA firing rates ^77,78^. The HD vectors were binned at 30°bins, ranging 0-360°for yaw and −60-60 for roll and pitch. The models were run *N* = 50 times each, with a random selection of *N* = 100 instances (or ‘trials’) of neural activity and HD at each HD bin. These models varied in the fraction of neurons used for fitting/testing, with 20,40,60,80, or 100% of neurons used. As in the Ridge models, the Logistic models also included varying time windows of neural activity used to decode any given time-point of the HD; these windows were 0.01, 0.05, 0.1, 0.2, 0.5, or 1.0 seconds.

### Lesions

Lesions of M2 were performed as previously described^23^, using excitotoxic injections of ibotenic acid (IA) (Abcam ab120041) delivered using an UMP3 UltraMicroPump (WPI) during two separate procedures. Aliquots of IA were prepared at 1% concentration and frozen. In the first procedure, IA was injected into four sites in one hemisphere (1.5 mm AP, relative to Bregma and 1.0 mm ML; 0.5 AP, 0.75 ML; −0.5 AP, 0.75 ML; and −1.5 AP, 0.75 ML, with two injections per site, at 1.6 and 0.8 mm below the brain surface, 75 nl each) and the animal was allowed to recover for one week, after which the injections were repeated at the same sites in the opposite hemisphere and electrode arrays were implanted in V1.

### Measuring Light Levels in Dark

To assess whether the recording box was sufficiently dark as to prevent the rats from being able to see anything in the dark recordings, we first made sure that it was impossible for a human observer (GG) to see any movement in the recording box after acclimating to the darkened room for 30 minutes. We then attempted to measure photon flux in the box using a photomultiplier tube (PMT) (ET Enterprises #9111B) after amplifying and filtering the signals (12dB lowpass at 10Hz) using a Stanford Research Systems preamplifier (#SR570).

Baseline PMT currents in the darkened behavioral box were measured with the PMT covered by tinfoil. After removing the tinfoil, the PMT current registered at 0.2*μ*A. This corresponds to 0.2 *10^−6^ Coulombs/s, which is 0.2 * 6.2415 *10^12^ electrons/s. Accounting for the PMT’s gain of 7.1 * 10^6^, that is 0.2 * 6.2415 * 10^12^*/*7.1 * 10^6^ or 1.8 * 10^5^ photocathode events/s. Given the PMT’s 10% quantum efficiency (QE) at 500nm, this corresponds to 1.8 * 10^6^ photons/s over the PMT’s 22mm cross-sectional area, or 80,000 photons/mm/s. Assuming 2.27*μm*^2^ rod cross section^79^, 0.4 specific absorption, and QE of 0.34^80^, that is 80000 * 2.27 * 10^−6^ or 0.18 incident photons/rod/s and finally 0.025 R*/rod/s. Based on retinal ganglion cell activity measured in Soucy *et al*.^81^, 0.025 R*/rod/s would correspond, roughly, to retinal ganglion cells firing at 6.7% of their peak firing rates measured at light levels corresponding to 100 R*/rod/sec.

### Statistics

All statistical comparisons were done using non-parametric tests (e.g. Mann-Whitney U or Wilcoxon) unless specified otherwise. A significance level of alpha = 0.01 was used throughout, unless otherwise noted. Bonferroni correction was applied where appropriate.

## Acknowledgements

We would like to thank Jeffrey Markowitz, Bence Ölveczky, and Sandeep Robert Datta, for advice and helpful discussions. Edward Soucy, Brett Graham, and Joel Greenwood of the CBS Neuroengineering Core were instrumentally helpful in technical advice and on light measurements. GG was supported by the National Science Foundation (NSF) Graduate Research Fellowship Program (GRFP).

## Author contributions statement

GG conceived and performed the electrophysiology and lesion experiments, analyzed the data, and wrote the manuscript. WL and JD provided assistance with modeling. DC provided research funding and space.

## Supplementary Figures

**Figure S1:**
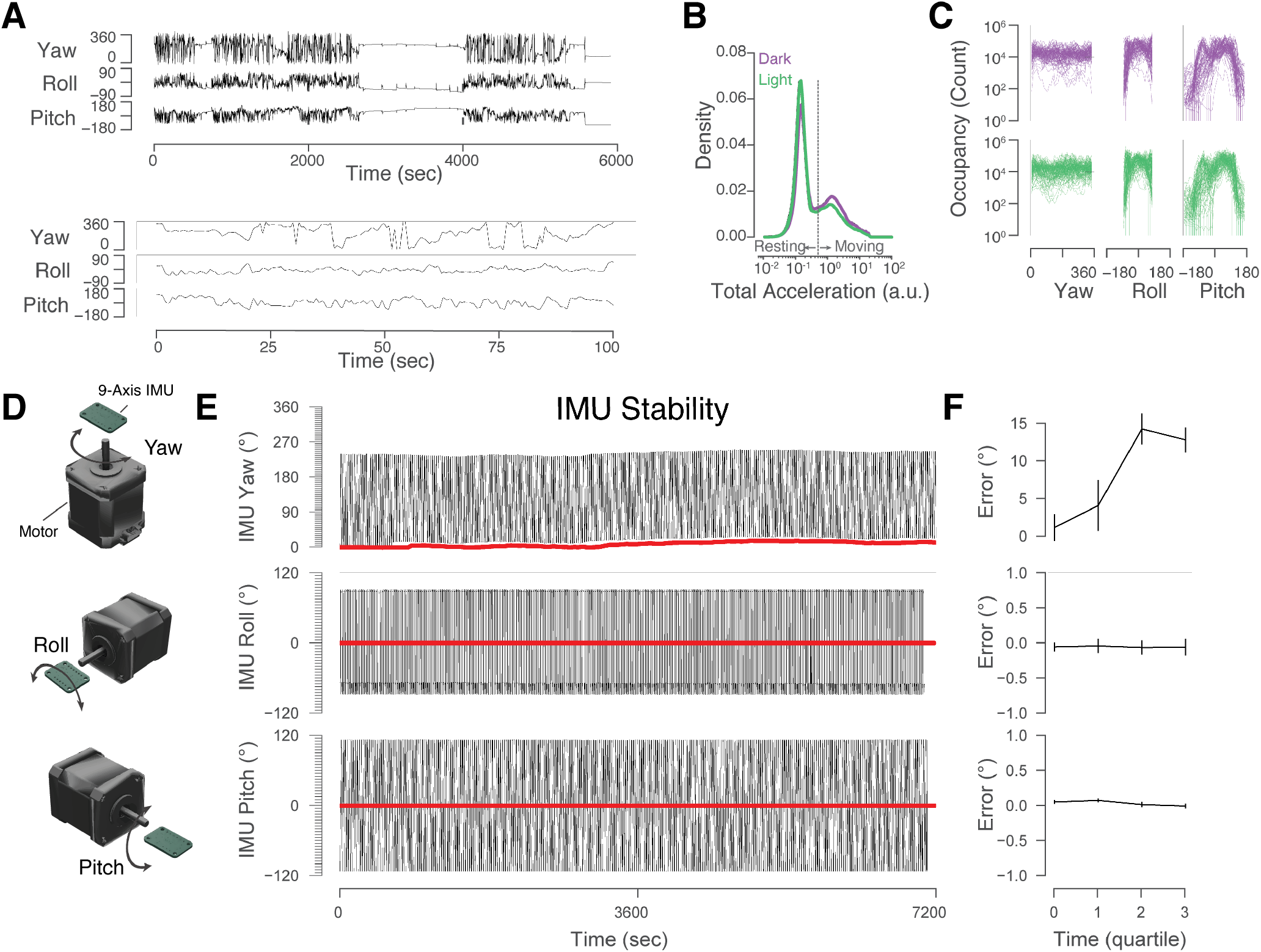
(Related to Figure 1). Details of Head Movement and IMU Stability. **(a)** Example traces from a recording session showing yaw, roll, and pitch components of head direction over 100 minutes (top) or 100 seconds (bottom). **(b)** Mean histograms of total acceleration, defined as the norm of the three linear components of acceleration, for sessions recorded in dark (purple) or light (green). **(c)** Behavioral occupancy for head direction angles split by individual two-hour sessions (each line). **(d)** Schematic of IMU stability test. The IMU PCB was attached to a stepper motor in one of three configurations to simulate movement along the yaw, roll, or pitch axes. A microcontroller rotated the motor back and forth for two hours while recording the resulting IMU signals. **(e)** Example traces of the stability test recordings showing yaw (top), roll (middle), and pitch (bottom) traces over time (black lines) and the accumulated measured angles (red). **(f)** Errors in the measured angle at quartiles of the two hour session relative to the beginning of the session.

**Figure S2:**
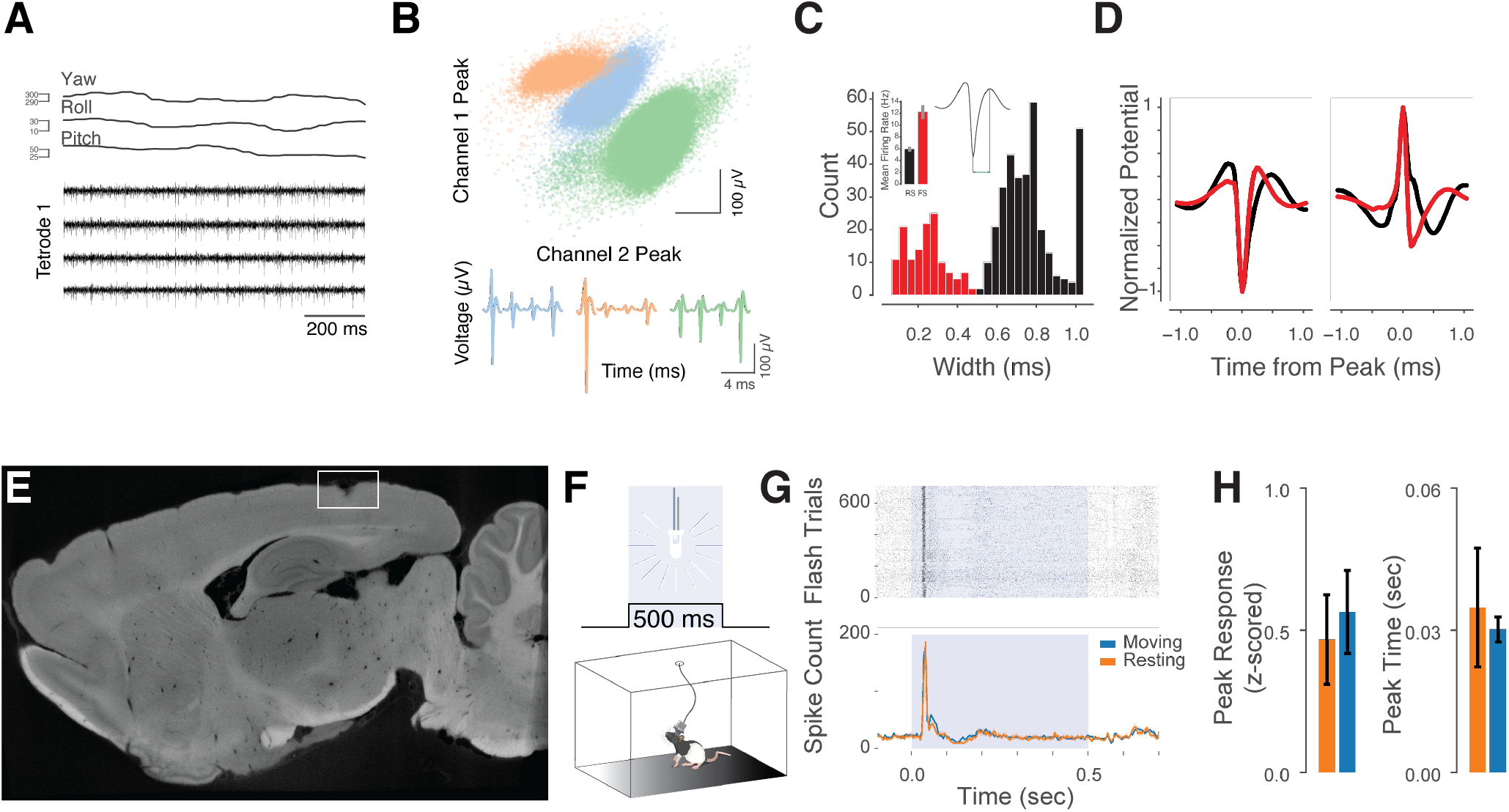
(Related to Figure 1). Details of V1 Neural Recordings. **(a)** Example traces from a recording session showing yaw, roll, and pitch components of head direction along with raw filtered neural activity traces from one example tetrode. **(b)** Spikes extracted from tetrodes were sorted into individual units based on waveform properties using Mountainsort ^74^. Top: example peak spike amplitudes from two channels on one tetrode. Colors indicate individual clusters from the sorting algorithm. Bottom: mean waveforms from each of the clusters, showing waveforms from each channel within the example tetrode. **(c)** Units were classified into regular-spiking putative pyramidal units (RSUs) or fast-spiking putative inhibitory units (FSUs) based on calculations of spike width (trough to peak time). For classification, waveforms from the highest-amplitude channels within a tetrode were used. Insets: bar graph of mean firing rates showing higher rates in FSUs than in RSUs (Red: FSUs (*N* = 61 in dark; *N* = 50 in light), Black: RSUs (*N* = 142 in dark; *N* = 112 in light)); and example waveform demonstrating width calculation. **(d)** Mean waveforms of *N* = 365 single units recorded in dark (*N* = 203) or light (*N* = 162) in *N* = 5 rats across *N* = 10 sessions, separated by negative-peaked (left) and positive-peaked (right) units. Color indicates fast-spiking units (FSUs, red) and regular-spiking units (RSUs, black) based on classification in **c. (e)** Sagittal view of a digital micro-CT image of a brain implanted with tetrodes in V1. White box outlines electrode track over V1. **(f)** Schematic of experiment to assess visual responsiveness of recorded neurons. Overhead LED lights were flashed (500-ms on, 400-600-ms random uniform off) while rats were freely moving in the recording chamber. **(g)** Example multiunit spike raster (top) and histogram (bottom) during flashes showing prominent responses to the visual stimulation. In the histogram, trials were separated by the animal’s movement status (resting or moving) at the time the LEDs turned on. **(h)** Summary of response magnitudes (left) and timing (right) to flashes across rats and sessions during movement or rest.

**Figure S3:**
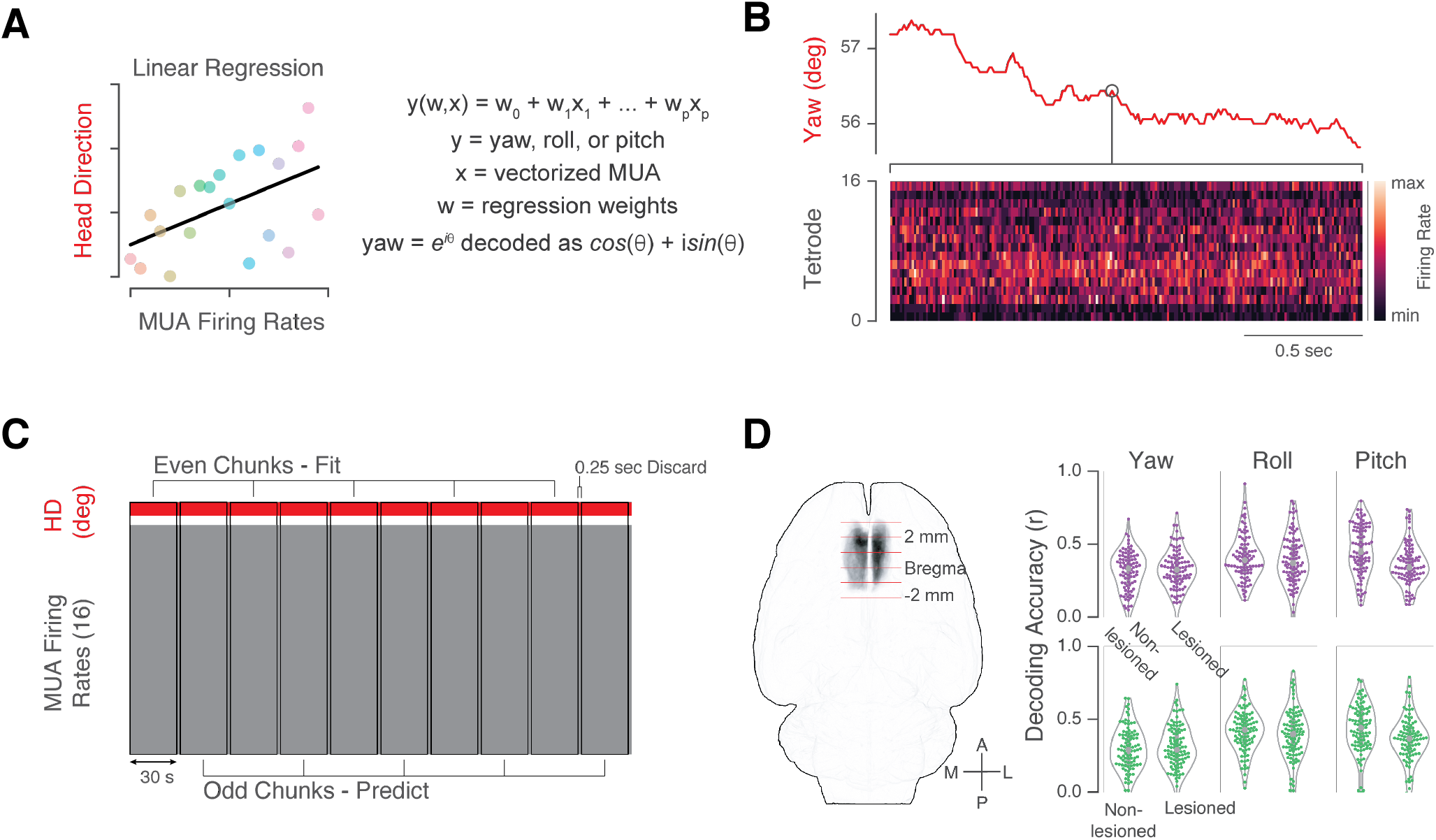
(Related to Figure 2). Head Direction Decoding Approach and Decoding in M2-Lesioned Animals. **(a)** Multiunit activity (MUA) vectors from the 16 tetrodes were used to predict yaw, roll, or pitch components of 3D HD separately, using ridge regression. **(b)** A vectorized MUA window ranging from 10 ms. to 2 sec. was used to predict any given HD time-point. **(c)** Sessions were divided into 30-second even and odd chunks, with the former used for fitting the models and the latter for testing. The chunks were separated by 0.25-second discarded gaps to ensure that model performance was not due to continuities in the HD or MUA signals. **(d)** Excitotoxic lesions to secondary motor cortex (M2) were performed bilaterally using ibotenic acid. Left: horizontal view of overlaid lesion ROIs from *N* = 4 rats. Right: Ridge regression decoding results for non-lesioned (*N* = 5) and lesioned (*N* = 4) rats. Each dot represents one 2-hour session; outlines are violin plots. Yaw, dark: Non-lesioned: *r*_*circ*_ = 0.31 ± 0.02; Lesioned: 0.33 ± 0.01 (mean ± SEM). MWU test *p* = 0.4. Yaw, light: Non-lesioned: *r*_*circ*_ = 0.30 ± 0.02; Lesioned: 0.31 ± 0.01. MWU test *p* = 0.31. Roll, dark: Non-lesioned: *r* = 0.41 ± 0.02; Lesioned: 0.39 ± 0.02. MWU test *p* = 0.26. Roll, light: Non-lesioned: *r* = 0.42 ± 0.01; Lesioned: 0.39 ± 0.02. MWU test *p* = 0.14. Pitch, dark: Non-lesioned: *r* = 0.46 ± 0.02; Lesioned: 0.36 ± 0.01. MWU test *p* = 1.2 × 10^−5^. Pitch, light: Non-lesioned: *r* = 0.44 ± 0.02; Lesioned: 0.36 ± 0.02. MWU test *p* = 7.5 × 10^−4^.

**Figure S4:**
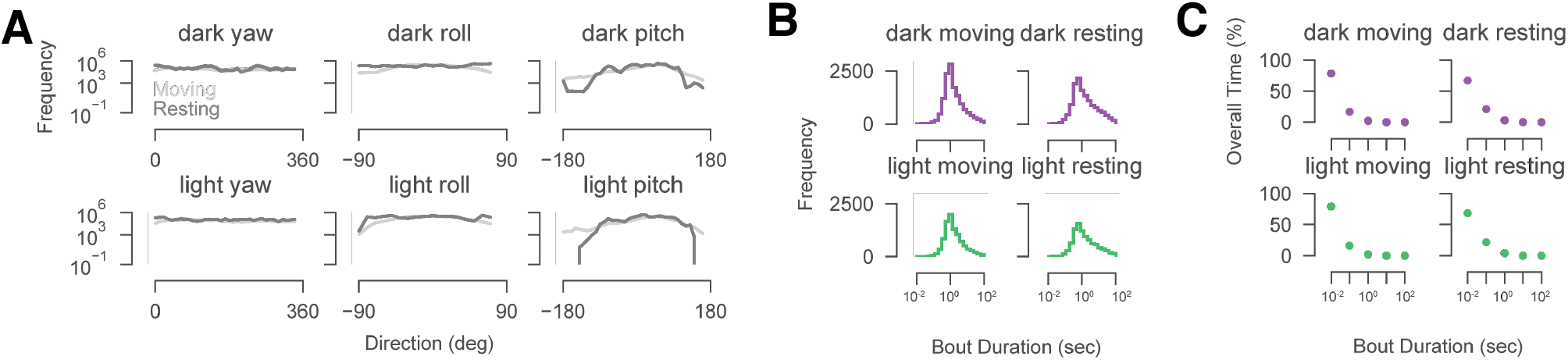
(Related to Figure 4). Behavioral Statistics During Movement and Rest. **(a)** Behavioral occupancy of binned yaw, roll, and pitch head directions split by moving (light gray) or resting (dark grey). **(b)** Distributions of durations of movement or rest bouts in dark or light sessions. Dark moving median: 1.18 seconds (6.84 ± 25.90 (mean ± std)); dark resting median: 1.09 seconds (9.00 ± 35.40). Light moving median: 1.21 seconds (6.20 ± 21.87); light resting median: 1.12 seconds (10.48 ± 38.31). **(c)** Overall time spent at or above various movement or rest bout durations.

**Figure S5:**
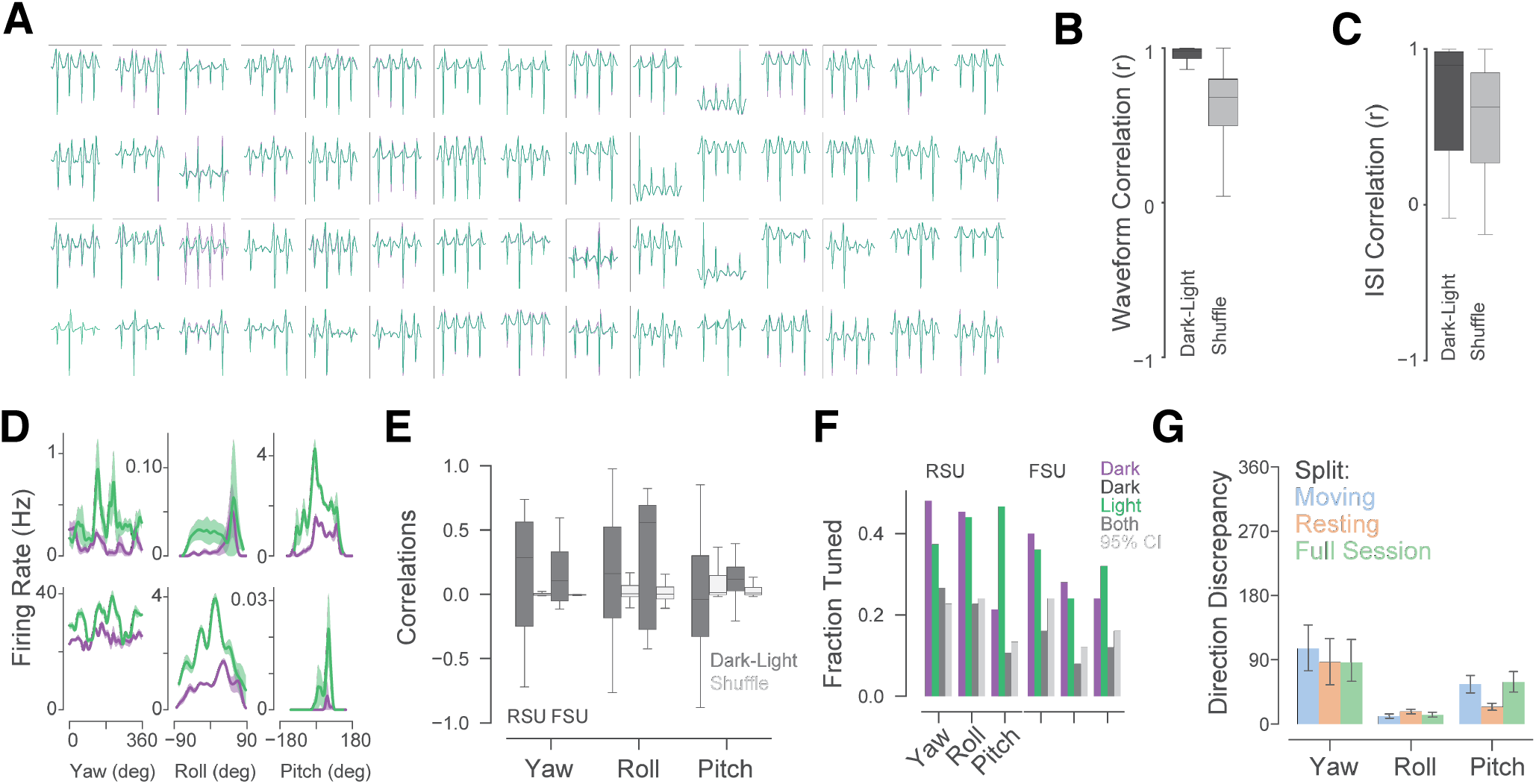
(Related to Figure 5). Tracking Single Units Across Dark and Light Sessions. **(a)** Example mean waveforms from tetrode channels for units sorted jointly across dark and light sessions (60 out of 100 units). **(b)** Correlations between mean waveforms in dark and light (*r* = 0.93±0.12, mean ± SEM) were significantly higher than among *N* = 1000 shuffled pairs (*r* = 0.55 ± 0.41), *p* = 2.57 × 10^−58^, MWU test. **(c)** Correlations between mean inter-spike-interval distributions for units recorded in dark and light (*r* = 0.70 ±0.34) were significantly higher than those among *N* = 1000 shuffled pairs (*r* = 0.56 ±0.32), *p* = 4.36 × 10^−09^, MWU test. **(d)** Example tuning curves of neurons recorded simultaneously in dark (purple) and light (green). **(e)** Correlations between tuning curves of neurons significantly tuned to HD in dark and light (dark grey), split by RSUs and FSUs; light gray: correlations of shuffled curves. **(f)** Fractions of neurons tuned to HD in dark (purple), light (green), or both conditions (dark grey). Fractions of neurons tuned in both light and dark were similar to the top 95% CI of a shuffled distribution (light grey). **(g)** Discrepancies in preferred directions between dark and light of neurons significantly tuned in both conditions.

## References

1 Bellmund, J. L., Gärdenfors, P., Moser, E. I. & Doeller, C. F. Navigating cognition: Spatial codes for human thinking. Science 362, eaat6766 (2018).

2 Buzsáki, G. & Moser, E. I. Memory, navigation and theta rhythm in the hippocampal-entorhinal system. Nature neuroscience 16, 130 (2013).

3 Kim, S. S., Hermundstad, A. M., Romani, S., Abbott, L. & Jayaraman, V. Generation of stable heading representations in diverse visual scenes. Nature 576, 126–131 (2019).

4 Kim, S. S., Rouault, H., Druckmann, S. & Jayaraman, V. Ring attractor dynamics in the drosophila central brain. Science 356, 849–853 (2017).

5 Seelig, J. D. & Jayaraman, V. Neural dynamics for landmark orientation and angular path integration. Nature 521, 186–191 (2015).

6 Fisher, Y. E., Lu, J., D’Alessandro, I. & Wilson, R. I. Sensorimotor experience remaps visual input to a heading-direction network. Nature 576, 121–125 (2019).

7 Nau, M., Schröder, T. N., Bellmund, J. L. & Doeller, C. F. Hexadirectional coding of visual space in human entorhinal cortex. Nature neuroscience 21, 188–190 (2018).

8 Nau, M., Julian, J. B. & Doeller, C. F. How the brain’s navigation system shapes our visual experience. Trends in cognitive sciences 22, 810–825 (2018).

9 Nau, M., Schroder, T. N., Frey, M. & Doeller, C. F. Behavior-dependent directional tuning in the human visual-navigation network. bioRxiv 765800 (2019).

10 Clark, B. J. & Taube, J. S. Vestibular and attractor network basis of the head direction cell signal in subcortical circuits. Frontiers in neural circuits 6, 7 (2012).

11 Yoder, R. M., Clark, B. J. & Taube, J. S. Origins of landmark encoding in the brain. Trends in neurosciences 34, 561–571 (2011).

12 Cullen, K. E. & Taube, J. S. Our sense of direction: progress, controversies and challenges. Nature Neuroscience 20, 1465 (2017).

13 Yoder, R. M. & Taube, J. S. The vestibular contribution to the head direction signal and navigation. Frontiers in integrative neuroscience 8, 32 (2014).

14 Taube, J. S. The head direction signal: origins and sensory-motor integration. Annu. Rev. Neurosci. 30, 181–207 (2007).

15 Angelaki, D. E. & Laurens, J. The head direction cell network: attractor dynamics, integration within the navigation system, and three-dimensional properties. Current Opinion in Neurobiology 60, 136–144 (2020).

16 Laurens, J. & Angelaki, D. E. The brain compass: a perspective on how self-motion updates the head direction cell attractor. Neuron 97, 275–289 (2018).

17 DiCarlo, J. J., Zoccolan, D. & Rust, N. C. How does the brain solve visual object recognition? Neuron 73, 415–434 (2012).

18 Niell, C. M. & Stryker, M. P. Modulation of Visual Responses by Behavioral State in Mouse Visual Cortex. Neuron 65, 472–479 (2010).

19 Leinweber, M., Ward, D. R., Sobczak, J. M., Attinger, A. & Keller, G. B. A Sensorimotor Circuit in Mouse Cortex for Visual Flow Predictions. Neuron 95, 1420–1432.e5 (2017).

20 Vélez-Fort, M. et al. A Circuit for Integration of Head-and Visual-Motion Signals in Layer 6 of Mouse Primary Visual Cortex. Neuron 1–25 (2018).

21 Gilbert, C. D. & Li, W. Top-down influences on visual processing. Nature Reviews Neuroscience 14, 350–363 (2013).

22 Saleem, A. B., Ayaz, A., Jeffery, K. J., Harris, K. D. & Carandini, M. Integration of visual motion and locomotion in mouse visual cortex. Nature Neuroscience 16, 1864–1869 (2013).

23 Guitchounts, G., Masís, J., Wolff, S. B. E. & Cox, D. Encoding of 3D Head Orienting Movements in the Primary Visual Cortex. Neuron 1–27 (2020).

24 Bouvier, G., Senzai, Y. & Scanziani, M. Head movements control the activity of primary visual cortex in a luminance-dependent manner. Neuron 108, 500–511 (2020).

25 Ji, D. & Wilson, M. A. Coordinated memory replay in the visual cortex and hippocampus during sleep. Nature neuroscience 10, 100–107 (2007).

26 Haggerty, D. C. & Ji, D. Activities of visual cortical and hippocampal neurons co-fluctuate in freely moving rats during spatial behavior. Elife 4, e08902 (2015).

27 Fiser, A. et al. Experience-dependent spatial expectations in mouse visual cortex. Nature Neuroscience 19, 1658–1664 (2016).

28 Saleem, A. B., Diamanti, E. M., Fournier, J., Harris, K. D. & Carandini, M. Coherent encoding of subjective spatial position in visual cortex and hippocampus. Nature 7, 1–18 (2018).

29 Keller, G. B. & Mrsic-Flogel, T. D. Predictive Processing: A Canonical Cortical Computation. Neuron 100, 424–435 (2018).

30 Chen, L. L., Lin, L.-H., Green, E. J., Barnes, C. A. & McNaughton, B. L. Head-direction cells in the rat posterior cortex. Experimental brain research 101, 8–23 (1994).

31 Chen, L. L., Lin, L. H., Barnes, C. A. & McNaughton, B. L. Head-direction cells in the rat posterior cortex. ii. contributions of visual and ideothetic information to the directional firing. Experimental brain research 101, 24–34 (1994).

32 Jacob, P.-Y. et al. An independent, landmark-dominated head-direction signal in dysgranular retrosplenial cortex. Nature neuroscience 20, 173 (2017).

33 Wallace, D. J. et al. Rats maintain an overhead binocular field at the expense of constant fusion. Nature 498, 65 (2013).

34 Meister, M. & Cox, D. Rats maintain a binocular field centered on the horizon. F1000Research 2 (2013).

35 Finkelstein, A. et al. Three-dimensional head-direction coding in the bat brain. Nature 517, 159–164 (2015).

36 Tukker, J. J., Tang, Q., Burgalossi, A. & Brecht, M. Head-directional tuning and theta modulation of anatomically identified neurons in the presubiculum. Journal of Neuroscience 35, 15391–15395 (2015).

37 Shinder, M. E. & Taube, J. S. Three-dimensional tuning of head direction cells in rats. Journal of neurophysiology 121, 4–37 (2019).

38 Xu, Z. et al. A comparison of neural decoding methods and population coding across thalamo-cortical head direction cells. Frontiers in neural circuits 13, 75 (2019).

39 Buzsáki, G. & Mizuseki, K. The log-dynamic brain: how skewed distributions affect network operations. Nature Reviews Neuroscience 15, 264–278 (2014).

40 Stackman, R. W. & Taube, J. S. Firing properties of rat lateral mammillary single units: head direction, head pitch, and angular head velocity. Journal of Neuroscience 18, 9020–9037 (1998).

41 Taube, J. S., Muller, R. U. & Ranck, J. B. Head-direction cells recorded from the postsubiculum in freely moving rats. i. description and quantitative analysis. Journal of Neuroscience 10, 420–435 (1990).

42 Super, H. & Roelfsema, P. R. Chronic multiunit recordings in behaving animals: advantages and limitations. Progress in brain research 147, 263–282 (2005).

43 Meyers, E. & Kreiman, G. Tutorial on pattern classification in cell recording. Visual population codes 517–538 (2012).

44 Schneider, D. M., Nelson, A. & Mooney, R. A synaptic and circuit basis for corollary discharge in the auditory cortex. Nature 513, 189–194 (2014).

45 Hengen, K. B., Pacheco, A. T., McGregor, J. N., Van Hooser, S. D. & Turrigiano, G. G. Neuronal Firing Rate Homeostasis Is Inhibited by Sleep and Promoted by Wake. Cell 165, 1–28 (2016).

46 Watson, B. O., Levenstein, D., Greene, J. P., Gelinas, J. N. & Buzsáki, G. Network homeostasis and state dynamics of neocortical sleep. Neuron 90, 839–852 (2016).

47 Timo-Iaria, C. et al. Phases and states of sleep in the rat. Physiology & behavior 5, 1057–1062 (1970).

48 Goodridge, J. P., Dudchenko, P. A., Worboys, K. A., Golob, E. J. & Taube, J. S. Cue control and head direction cells. Behavioral neuroscience 112, 749 (1998).

49 Calton, J. L. & Taube, J. S. Degradation of head direction cell activity during inverted locomotion. Journal of Neuroscience 25, 2420–2428 (2005).

50 Angelaki, D. E. et al. A gravity-based three-dimensional compass in the mouse brain. Nature communications 11, 1–13 (2020).

51 Yartsev, M. M. & Ulanovsky, N. Representation of three-dimensional space in the hippocampus of flying bats. Science 340, 367–372 (2013).

52 Grieves, R. M. et al. The place-cell representation of volumetric space in rats. Nature communications 11, 1–13 (2020).

53 Page, H. J., Wilson, J. J. & Jeffery, K. J. A dual-axis rotation rule for updating the head direction cell reference frame during movement in three dimensions. Journal of neurophysiology 119, 192–208 (2018).

54 LaChance, P. A., Dumont, J. R., Ozel, P., Marcroft, J. L. & Taube, J. S. Commutative properties of head direction cells during locomotion in 3d: are all routes equal? Journal of Neuroscience 40, 3035–3051 (2020).

55 Taube, J. S., Wang, S. S., Kim, S. Y. & Frohardt, R. J. Updating of the spatial reference frame of head direction cells in response to locomotion in the vertical plane. Journal of Neurophysiology 109, 873–888 (2013).

56 M-lynarski, W. F. & Hermundstad, A. M. Adaptive coding for dynamic sensory inference. Elife 7, e32055 (2018).

57 Ganguli, D. & Simoncelli, E. P. Efficient sensory encoding and bayesian inference with heterogeneous neural populations. Neural computation 26, 2103–2134 (2014).

58 Heeger, D. J. Theory of cortical function. Proceedings of the National Academy of Sciences 114, 1773–1782 (2017).

59 Schor, R. H., Miller, A. D. & Tomko, D. L. Responses to head tilt in cat central vestibular neurons. i. direction of maximum sensitivity. Journal of Neurophysiology 51, 136–146 (1984).

60 Mimica, B., Dunn, B. A., Tombaz, T., Bojja, V. P. T. N. C. S. & Whitlock, J. R. Efficient cortical coding of 3D posture in freely behaving rats. Science 362, 584–589 (2018).

61 Taube, J. S. & Burton, H. L. Head direction cell activity monitored in a novel environment and during a cue conflict situation. Journal of Neurophysiology 74, 1953–1971 (1995).

62 Zong, W. et al. Large-scale two-photon calcium imaging in freely moving mice. bioRxiv (2021).

63 Skaggs, W. E., Knierim, J. J., Kudrimoti, H. S. & McNaughton, B. L. A model of the neural basis of the rat’s sense of direction. In Advances in neural information processing systems, 173–180 (1995).

64 Rao, R. P. & Ballard, D. H. Predictive coding in the visual cortex: a functional interpretation of some extra-classical receptive-field effects. Nature Neuroscience 2, 79–87 (1999).

65 Clark, A. Whatever next? predictive brains, situated agents, and the future of cognitive science. Behavioral and brain sciences 36, 181–204 (2013).

66 Friston, K. A theory of cortical responses. Philosophical transactions of the Royal Society B: Biological sciences 360, 815–836 (2005).

67 Siegle, J. H. et al. Open Ephys: an open-source, plugin-based platform for multichannel electrophysiology. Journal of Neural Engineering 14, 045003–14 (2017).

68 Nguyen, D. P. et al. Micro-drive array for chronic in vivo recording: tetrode assembly. JoVE (Journal of Visualized Experiments) e1098 (2009).

69 Kloosterman, F. et al. Micro-drive array for chronic in vivo recording: drive fabrication. JoVE (Journal of Visualized Experiments) e1094 (2009).

70 Mendoza, G. et al. Recording extracellular neural activity in the behaving monkey using a semi-chronic and high-density electrode system. Journal of Neurophysiology 116, jn.00116.2016–574 (2016).

71 Vandecasteele, M. et al. Large-scale recording of neurons by movable silicon probes in behaving rodents. JoVE (Journal of Visualized Experiments) e3568 (2012).

72 Ferguson, J. E., Boldt, C. & Redish, A. D. Creating low-impedance tetrodes by electroplating with additives. Sensors and actuators. A, Physical 156, 388–393 (2009).

73 Dhawale, A. K. et al. Automated long-term recording and analysis of neural activity in behaving animals. eLife 6, 91 (2017).

74 Chung, J. E. et al. A Fully Automated Approach to Spike Sorting. Neuron 95, 1381–1394.e6 (2017).

75 Giocomo, L. M. et al. Topography of head direction cells in medial entorhinal cortex. Current Biology 24, 252–262 (2014).

76 Pedregosa, F. et al. Scikit-learn: Machine learning in python. Journal of Machine Learning Research 12, 2825–2830 (2011).

77 Hung, C. P., Kreiman, G., Poggio, T. & DiCarlo, J. J. Fast readout of object identity from macaque inferior temporal cortex. Science 310, 863–866 (2005).

78 Georgopoulos, A. P., Schwartz, A. B. & Kettner, R. E. Neuronal population coding of movement direction. Science 233, 1416–1419 (1986).

79 Hagins, W., Penn, R. & Yoshikami, S. Dark current and photocurrent in retinal rods. Biophysical journal 10, 380–412 (1970).

80 Nymark, S., Heikkinen, H., Haldin, C., Donner, K. & Koskelainen, A. Light responses and light adaptation in rat retinal rods at different temperatures. The Journal of physiology 567, 923–938 (2005).

81 Soucy, E., Wang, Y., Nirenberg, S., Nathans, J. & Meister, M. A novel signaling pathway from rod photoreceptors to ganglion cells in mammalian retina. Neuron 21, 481–493 (1998).

